# Longitudinal investigation of prostate tumor spheroid proliferation with dynamic line-field optical coherence tomography

**DOI:** 10.1101/2025.11.04.686594

**Authors:** Steph Swanson, Keyu Chen, Elahe Cheraghi, Ernest Osei, Kostadinka Bizheva

## Abstract

Recently, it has become widely recognized that culturing cancer cells *in vitro* in small, 3D aggregates known as tumor spheroids provides a more physiologically relevant model of *in vivo* tumor behavior compared to 2D monolayer cultures. Dynamic optical coherence tomography (dOCT) is a non-invasive imaging modality that, by analyzing temporal fluctuations in the light scattered from biological tissue, does not require exogenous contrast agents to visualize and quantify cellular activity within 3D cell cultures. However, recent volumetric dOCT studies have encountered challenges due to low acquisition speeds. In this study, we present morphological and dynamic analyses of prostate tumor spheroid growth over a two-week longitudinal period, utilizing volumetric imaging with a line-field dOCT platform. Our method clearly differentiated between active cellular metabolism in live spheroids and the lack of activity in spheroids fixed with formaldehyde. Quantitative validation of the dynamic signal was conducted using the Alamar Blue proliferation assay, while qualitative validation was provided by live/dead fluorescence microscopy.

## 1 Introduction

Prostate cancer is the most commonly diagnosed cancer among North American men [1, 2]. The use of *in vitro* cell culture has been invaluable in the investigation of tumor cell biology and cancer progression [3]. Historically, tumors were thought to be composed of a homogeneous population of cancer cells [4], which was a perspective well-suited to the conventional practice of culturing cells *in vitro* as 2D monolayers on flat, rigid surfaces. In a 2D monolayer, each cell has equal and unrestricted access to oxygen, nutrients, and signaling molecules available in the culture medium. However, cancer cell behavior is influenced by its surrounding environment [5], including nearby cells [6], signaling molecules [4], and structural features such as the extracellular matrix (ECM) [7] and cell adhesion molecules like integrin [8]. Although 2D cell culture methods are efficient, scalable, and compatible with many biological assays, cells grown as monolayers lack physiologically relevant cell-cell and cell-environment interactions [9]. These interactions critically influence cellular signaling, proliferation, and gene and protein expression [10, 11]. In recent decades, it has been demonstrated that culturing cancer cells as small, 3D aggregates, known as tumor spheroids, provides improved physiological relevance [12, 13]. The increased cell-to-cell contact and spatial organization within *in vitro* tumor spheroids promote cellular morphologies and characteristics that more closely resemble those found in *in vivo* tumors [14, 15]. Unlike the homogeneous environment in monolayer cultures, diffusion-limited nutrient distribution in tumor spheroids gives rise to heterogeneous cell populations: proliferative cells form the outer layer in contact with the culture medium, whereas the inner core accumulates metabolic waste and contains necrotic and apoptotic cells [16].

Nevertheless, the 3D nature of tumor spheroids challenges conventional biological tools, which typically destroy spatial information either by averaging signals across a population of cells or by requiring disaggregation of cellular structures into cell suspensions. For example, a common method of assessing cellular activity is the non-toxic Alamar Blue (AB) proliferation assay, which measures cellular metabolism through the reduction of the blue, non-fluorescent dye resazurin to the pink, highly fluorescent resorufin [17]. However, the fluorescence measured in this assay directly quantifies the total amount of resorufin produced within the spheroid, providing only an average metric [18]. Another widely used method for spheroid assessment is fluorescence microscopy (FM). Although this modality preserves spatial information and does not require spheroid disaggregation, fluorescence measurements remain qualitative rather than quantitative. Additionally, introducing fluorophores into the sample is invasive and precludes true longitudinal investigation.

Optical coherence tomography (OCT) is a non-invasive optical imaging modality that relies on low-coherence interferometry to generate volumetric images of biological samples with micrometer-scale resolution [19, 20]. Previous studies have employed OCT to provide insights into the 3D structure of tumor spheroids [21–25]. The application of OCT has expanded beyond static morphological imaging to dynamic OCT (dOCT), which analyzes temporal fluctuations in repeated OCT acquisition [26]. Unlike contrast derived from exogenous fluorophores or stains, the contrast in dOCT images is generated non-invasively by quantitative statistical analysis of the intensity fluctuations of the optical beam registered at each pixel of the OCT detector. dOCT has enhanced investigations of various 3D cell culture models, including tumor spheroids [27–38], organoids [39–44], paper-based 3D culture [45], and 3D scaffolds [46]. Furthermore, dOCT has also been utilized for cancer diagnosis in human biopsy samples from lung [47], breast [48], and skin [49] tissues.

While OCT was initially used to study particle dynamics (Brownian motion) in microsphere suspensions [50, 51], the idea of using temporal fluctuations in the OCT signal generated by light scattered from single cells to study cell apoptosis was first proposed in 2010 by Van der Meer et al. [52]. Since then, numerous dOCT approaches have been developed for the different designs of high speed OCT technology: spectral domain OCT (SD-OCT) [27, 29, 36, 38, 39, 53–58], swept source OCT (SS-OCT) [30–33, 59], full field OCT (FF-OCT) [28, 42, 43, 60–66], and line-field OCT (LF-OCT) [37]. Furthermore, various dOCT methods for analysis and interpretation of the OCT interferogram itself have been employed, including a standard-deviation metric termed motility [27, 39], running standard deviation [60], autocorrelation [28, 39, 53], OCT correlation decay speed (OCDS) [30], intensity variance [29], logarithmic intensity variance (LIV) [30] and its modifications [59], and median adjusted variance (MAV) [36]. Other methods have characterized the spectrum of the OCT interferogram by fitting the power spectral density (PSD) to an inverse-power-law model [39], integrating it over discrete frequency bands to generate red-green-blue (RGB) images [37, 54–58, 61, 62], or computing the mean frequency and inverse frequency bandwidth to generate hue-saturation-brightness (HSB) images [38, 42, 43, 63–66]. Many dOCT assessments of 3D cell cultures have been qualitatively compared with established biological techniques, including FM [30–33], immunohistochemistry [42], TUNEL assay [43], histological staining [44], and luciferase-based ATP luminescence assay and live/dead cell counting via nuclear staining [46]. However, direct and quantitative validation of the dOCT measurements against standard biological methods has been limited and exclusively performed with point-scanning (PSc) OCT. These studies analyzed fluctuations in the OCT intensity signal itself through motility and the inverse-power-law [40, 45] or MAV [36], or analyzed its spectrum via the HSB method [38] or the mean frequency alone [67]. Compared to FF-OCT systems, PSc-OCT offers an approximate 15 dB sensitivity gain [68] and faster dOCT volumetric acquisition (compared to time-domain FF-OCT) [54], which is required for the investigation of 3D cell culture. Nevertheless, extended data acquisition times have limited previous studies to cross-sectional imaging or temporally sparse volumetric data collection. To bridge this gap, we recently developed a high-speed spectral-domain LF-OCT system for volumetric dOCT imaging [37]. We adapted the frequency banding technique [61], which to the best of our knowledge, has never been quantitatively validated to an established biological method.

In this study, we employed the LF-dOCT technology and analytical method to longitudinally investigate qualitative and quantitative morphological and dynamic changes in prostate tumor spheroids at the cellular level over two weeks of growth.

## 2 Methods

### 2.1 OCT setup

In this study, we employed a high-speed spectral-domain LF-dOCT system [37] for both morphological and dynamic imaging of tumor spheroids. The system utilizes a broadband superluminescent diode (SLD) light source (cBLMD-T-850-HP, Superlum, Ireland) with a central wavelength of 842 nm and a full-width-at-half-maximum (FWHM) spectral bandwidth of 179 nm, resulting in an axial OCT resolution of 2.6 *µ*m in air, equivalent to 1.9 µm in biological tissue assuming an average refractive index of 1.38. A high-speed area CMOS camera (FASTCAM NOVA S9, Photron) is used at the detection end of the LF-OCT system to acquire B-scans at a maximum rate of 2,500 fps. Utilizing 1024 camera pixels in the spectral direction, the system achieves an axial scanning range of approximately 1 mm. With a microscope objective (M Plan APO, Mitutoyo, Japan) of 5x magnification power in the LF-OCT imaging probe, the system’s lateral resolution is 6.4 *µ*m in air. For an optical power of 3.5 mW incident on the imaged object and an imaging rate of 2,000 fps, the system’s sensitivity was approximately 93 dB. A detailed description of the system’s design and performance can be found in [37].

### 2.2 Cell culture methods

#### 2.2.1 Spheroid culture

PC3 cells were cultured in RPMI 1640 (Sigma-Aldrich, USA) supplemented with 10% fetal bovine serum (Thermo Fisher Scientific, USA) and 1% penicillin (Thermo Fisher Scientific, USA). Cells were maintained at 37°C with 5% CO_2_ in a humidified atmosphere. Prostate tumor spheroids were created and cultured in a 3D Petri Dish (MicroTissues Inc., USA) made of 2% agarose gel and containing a 9 × 9 array of wells that were each seeded with approximately 1,000 cells (Fig. 1A). Here, one 3D Petri Dish is referred to as a gel. Culture medium in each gel was exchanged every other day.

**Figure 1.**
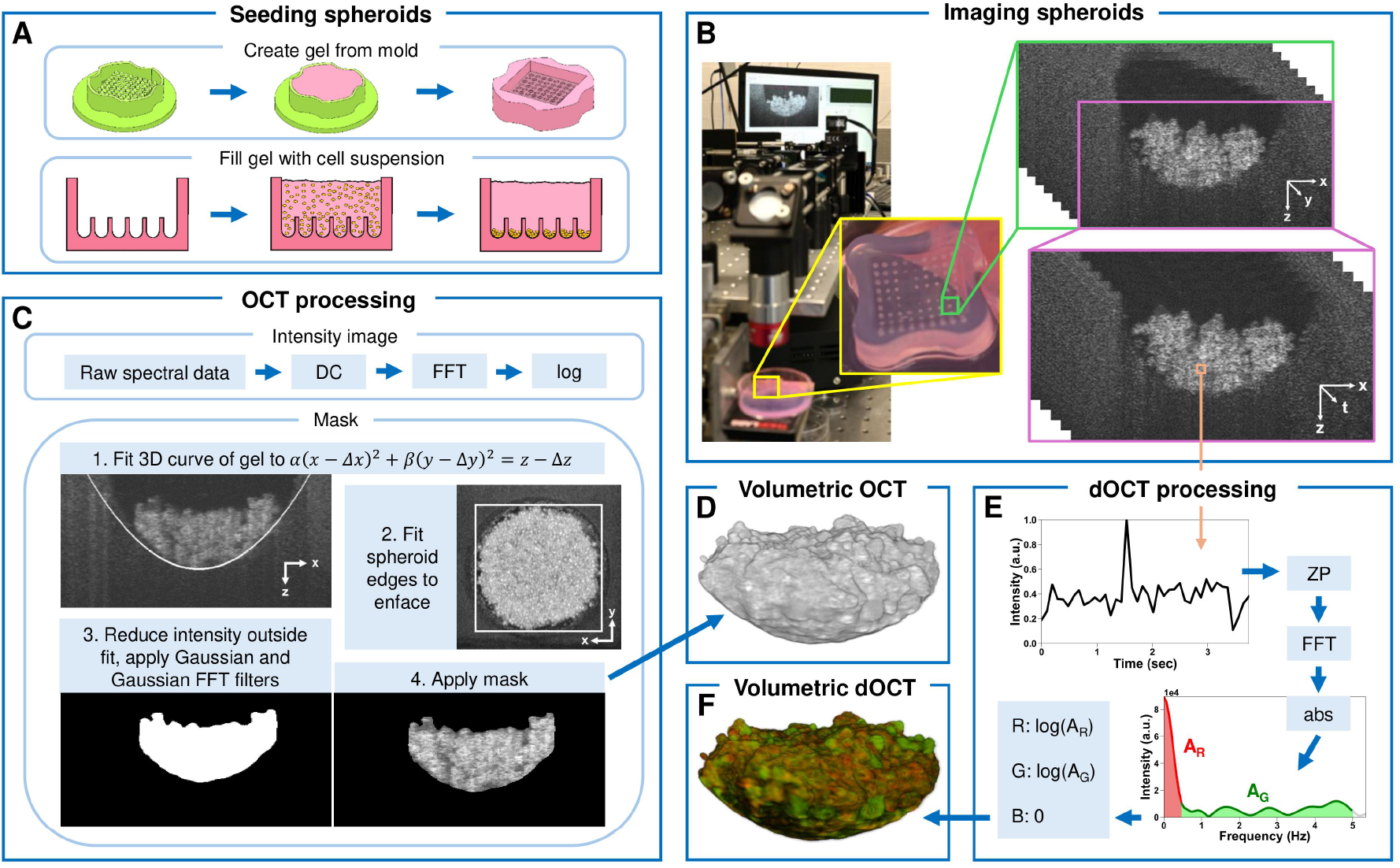
Schematic diagram of setup and processing. (A) Seeding spheroids. Agarose gels, depicted in pink, that each contained an array of wells were filled with a suspension of cells, shown in yellow, to form prostate tumor spheroids at the bottom of each well. (B) Imaging spheroids. Spectral-domain LF-dOCT system with glass Petri dish containing a gel of spheroids. The yellow inset magnifies the gel to show spheroids in the wells. The green inset displays a single spheroid in its well with a volumetric collection of B-scans. The purple inset displays the collection of B-scans acquired at one location over 4 seconds of recording time. (C) OCT processing. Intensity images were used to generate masks by reducing intensity below 3D curve of well inside gel (Eq. 1), beyond lateral edges of spheroid, and with Gaussian and Gaussian FFT filters. (D) Volumetric OCT image of live spheroid generated from mask. (E) dOCT processing at each voxel location by FFT in time direction. Integration of spectrum was performed along two frequency bands: red over 0-0.5 Hz and green over 0.5-5 Hz. (F) Volumetric dOCT image of live spheroid. DC: dispersion compensation; FFT: fast Fourier transform; log: base 10 logarithm; ZP: zero-padding; abs: absolute value; *A*_*R*_: area under red frequency band; *A*_*G*_: area under green frequency band.

#### 2.2.2 Pseudo-longitudinal protocol

OCT imaging of the cell seeding process began immediately after the cell suspension was introduced into the well. Without moving the imaging stage of the LF-OCT system, images were acquired regularly for two hours as cells settled to the bottom of the well. Longitudinal dOCT image acquisition of spheroid growth began 24 hours after seeding and was repeated every 24 hours for two weeks. During each imaging session, 15 live tumor spheroids were imaged, then fixed with 4% formaldehyde (Thermo Fisher Scientific, USA) in PBS (Wisent Inc., Canada) for 1 hour to remove metabolic activity and subsequently maintained at 4°C for ~24 hours to reduce motion induced during the fixing process. Afterward, 15 of the spheroids fixed during the previous day’s imaging session were also imaged. During imaging, spheroids were kept at 21°C without CO_2_ supply. As spheroids were sacrificed by fixation after each imaging session, 42 gels were seeded at the beginning of the experiment, with three gels allocated for each of the 14 days of imaging.

#### 2.2.3 Alamar Blue proliferation assay

Average cellular proliferation within the tumor spheroids was measured via AB proliferation assay. At each measurement time, culture medium was removed from three gels of spheroids and replaced with 34.5 *µ*L of AB (Sigma-Aldrich, USA) and 156 *µ*L of RPMI to achieve 5% concentration throughout each 500 *µ*L gel. Samples were then incubated for 3 hours at 37°C with 5% CO_2_ in a humidified atmosphere before intensity was recorded using a BioTek Synergy H1 microplate reader at 570 nm. Gels containing no cells were treated identically and measured as background. The average background intensity was subtracted from the measurement of gels containing spheroids. After background subtraction, AB absorbance was normalized relative to the absorbance measured on day 1.

#### 2.2.4 Live/dead fluorescence microscopy

FM was used to spatially assess cell death within spheroids using a Live/Dead Cell Viability assay (Sigma-Aldrich, USA) comprising two fluorophores. The first, calcein acetoxymethyl (calcein-AM), is a cell-permeant dye that identifies live cells through conversion to fluorescent green calcein by intracellular processes [69]. The second, propidium iodide (PI), is not permeant to live cells and thus instead identifies dead cells by fluorescing red upon binding with DNA or RNA [70]. Culture medium was removed from each gel and replaced with a solution of 0.2875 *µ*L calcein-AM, 1.15 *µ*L PI, 95 *µ*L RPMI, and 95 *µ*L PBS. Gels were incubated for one hour at 37°C with 5% CO_2_ in a humidified atmosphere before transfer to cover slip-bottomed Petri dishes that were placed on the stage of a Zeiss Axio Observer widefield microscope. The microscope was operated by ZEN2 Blue Edition and equipped with an Axiocam 506 mono camera. Fluorescent images were acquired with a FOV of 1.25 mm by 1 mm and a lateral resolution of 0.908 *µ*m using a Plan-Apochromat 63/1.4 Oil Ph3 M27 objective (Carl Zeiss Microscopy-LLC, NY, USA). Temperature was maintained at 37°C throughout imaging using an incubating plate (PECON, Erbach, Germany).

### 2.3 OCT Image acquisition

Both morphological and dynamic OCT images were acquired with the LF-dOCT system at a B-scan rate of 2,000 fps. Each B-scan frame consisted of 400 A-scans, and one 3D volume comprised 400 B-scans. For cell seeding, only morphological OCT data were acquired. At each time point, 5 repeated volumes were acquired consecutively and averaged to improve image contrast. For longitudinal dynamic imaging, each dOCT volume was divided into four sub-volumes, with 40 repetitions per sub-volume, corresponding to approximately four seconds of recording time, resulting in a sub-volume rate of 10.7 Hz. This sub-volume acquisition frequency was approximately half of the theoretical rate given the camera speed due to the non-linear behavior of the scanner, as described in detail in [37]. To address the scanner’s non-linearity, 20 additional frames were acquired and discarded at the ends of each sub-volume and the rest time of the scanner was equal to 50% of the acquisition time for a proper reset. The total dOCT recording time for one tumor spheroid was approximately 16 seconds.

### 2.4 Image processing

#### 2.4.1 Morphological

Morphological OCT images were generated using a standard OCT image reconstruction process. The resulting images were digitally dispersion-compensated and displayed on a logarithmic scale. In this work, we developed a novel masking protocol to isolate the spheroid from the surrounding gel (Fig. 1C). To generate the mask, the 3D curve of the well containing the spheroid within the gel was first fit to the curve:

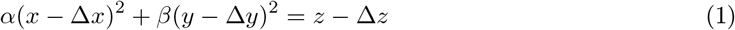

where *x* and *z* are respectively the lateral and depth positions in each B-scan, *y* is the position along the lateral scanning direction, and the location of the bottom of the well is given by (Δ*x*, Δ*y*, Δ*z*). The steepness of the well in the *x* and *y* directions is given by *α* and *β*, respectively. Then, the intensity was reduced below the 3D curve of the well and beyond a manually selected lateral rectangular region of interest (ROI) on the maximum intensity projection enface, chosen to contain the spheroid. Intensity far from the center of the spheroid was further reduced using a large Gaussian filter with FWHM comparable to the width of each B-scan. Finally, a Gaussian FFT filter with *σ* = 25 was applied, and a volumetric mask was generated by removing all pixels with zero intensity. In particularly dense spheroids, incident light did not penetrate well to the bottom, resulting in much lower intensity. In these cases, the masks were adjusted to include portions of the spheroid that the filters had removed.

Then, a newly developed numerical morphological analysis on the volumetric masks was carried out. The volume *V*_sph_ and external surface area *S* of each spheroid were estimated directly as the volume of non-zero pixels in the mask and the externally bounding surface area, respectively. The volume of internal gaps *V*_gap_ was estimated as the volume of zero intensity pixels within the external surface area. The surfaces of internal gaps were not included in the external surface area. To compare the external surface area of each spheroid to the volume it enclosed, the following sphericity metric was developed:

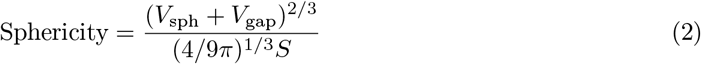

Since the ratio *V* ^2*/*3^*/S* of a solid sphere is (4*/*9*π*)^1/3^, sphericity achieves a maximum of 1 for any sphere, regardless of whether the sphere is hollow or full.

#### 2.4.2 Dynamic

The dOCT algorithm employed in this study investigates motion of cells in tumor spheroids using frequency banding [61] (Fig. 1E). To generate dOCT B-scans, a Hann window function was first applied to the linearly scaled OCT intensity values at each voxel location in the time dimension to reduce sidelobe artifacts and frequency leakage after FFT. Following zero-padding to improve digital resolution, FFT was applied along the time-axis at each location. For ease of comparison with live/dead FM, integrals along two frequency bands of interest were calculated: a red band of slow frequencies below 0.5 Hz and a green band of fast frequencies between 0.5 and 5 Hz. Logarithmic scaling was applied to each of the red and green dOCT channels to recalibrate the intensity distribution. To mitigate random variations in the recorded signal and background intensities, a uniform intensity baseline was achieved by subtracting the sum of the average and standard deviation of the intensity within a small region of culture medium above each spheroid in a central B-scan. Unlike in our previous work [37], to display the combined channels as a color image, each masked channel was independently normalized to those of a representative spheroid. Here, to reduce random noise and improve image quality, a Gaussian FFT filter with *σ*_*R*_ = 90 and *σ*_*G*_ = 65 to the red and green channels, respectively, was also applied, followed by the previously utilized 3 × 3 median filter.

In our newly developed numerical dynamic analysis, the above process was followed with the exception of normalization to a representative spheroid. A raw dynamic signal metric was developed for a given spheroid as:

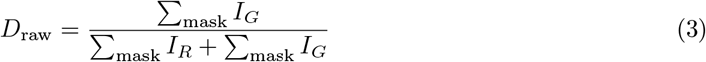

where *I*_*R*_ and *I*_*G*_ represent the red and green intensity channels, respectively, and ∑_mask_ is a voxel-wise sum over the masked spheroid. To account for the lower intensity in dense spheroids, the intensity above the 3D curve of the well and within the lateral ROI was used to exclude C-scans from numerical analysis whose average intensity was lower than the volume-wide median. The average raw dynamic signal of each day’s fixed spheroids was subtracted from that of the corresponding live spheroids, and the result was normalized relative to day 1 as:

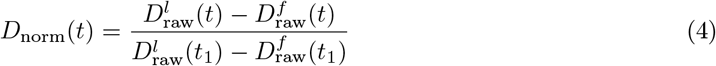

Here, 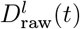 and 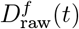 are respectively the live and fixed raw dynamic signal on the day given by *t*, with day 1 given by *t*_1_.

### 2.5 Statistics

All measurements are presented as averages with standard deviation. Numerical OCT and dOCT measurements that fell outside of the interquartile range were excluded.

## 3 Results

### 3.1 Morphological OCT

Figure 2 summarizes results from the morphological OCT imaging. The first row displays unmasked central B-scans of spheroid formation 2, 5, 10, 60, and 120 minutes after seeding. The culture medium inside the well appears dark, while the agarose gel exhibits faint speckles due to its relatively low reflectivity. Cells appear suspended in the culture medium at 2 minutes after seeding (Fig. 2A) but by 5 minutes they have settled at the bottom of the well (Fig. 2B). Subsequently, cells self-assemble into spheroids by forming clumps that condense and gradually pull away from the gel at 10, 60, and 120 minutes after seeding (Fig. 2C-E).

**Figure 2.**
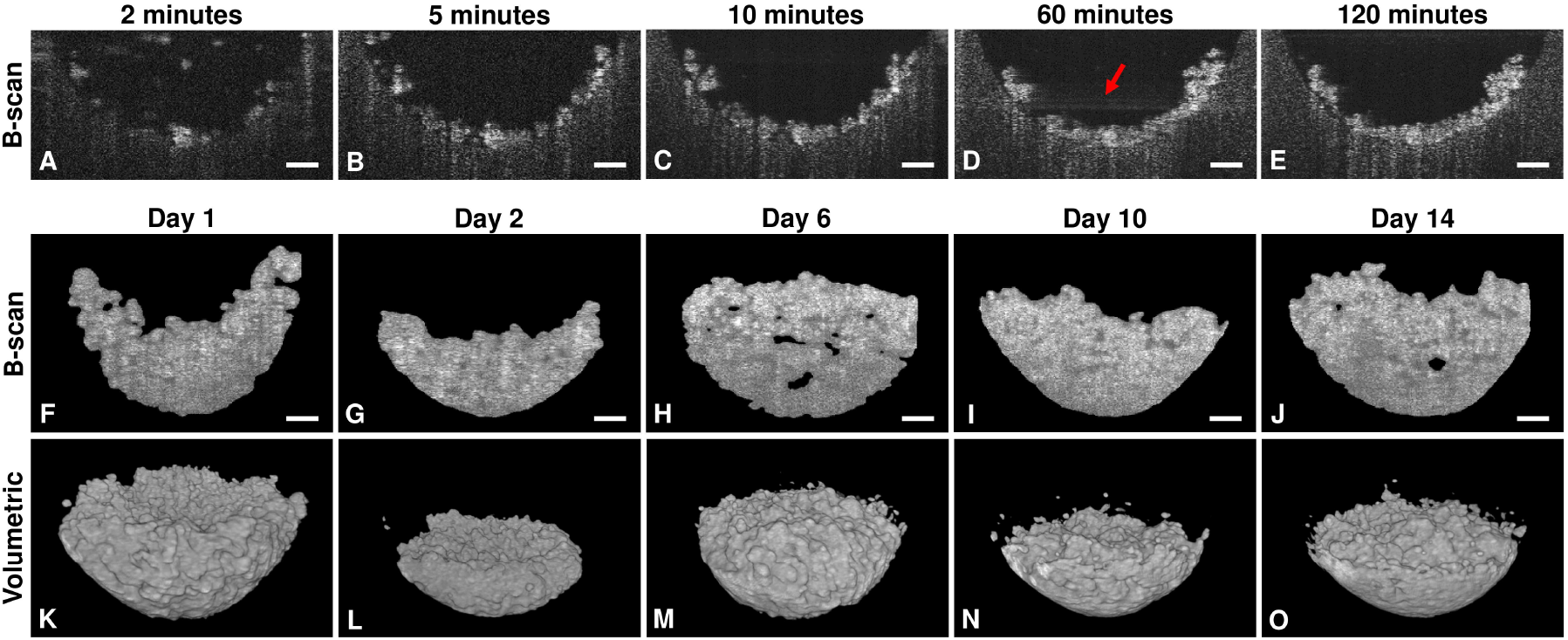
Morphological OCT images of prostate tumor spheroids. Central B-scan OCT images (A) 2, (B) 5, (C) 10, (D) 60, and (E) 120 minutes after cell seeding. Reflection artifact is labeled with red arrow. Masked central B-scan OCT images (F) 1, (G) 2, (H) 6, (I) 10, and (J) 14 days after cell seeding. Masked volumetric OCT images (M) 1, (N) 2, (O) 6, (P) 10, and (Q) 14 days after cell seeding. All scale bars are 100 *µ*m.

The second and third rows of Figure 2 show central B-scans and corresponding volumetric OCT images of representative spheroids on days 1, 2, 6, 10, and 14, respectively. The gel and culture medium surrounding each spheroid have been masked. Despite the spheroids appearing quite flat 120 minutes after seeding (Fig. 2E), as they grow, their upper portions became more spherical while the bottom portion retained the parabolic shape of their containing well. On day 1, the central region of the spheroids appeared collapsed (Fig. 2F) whereas the periphery grew diffusely along the slope of the well (Fig. 2K), similar to the cellular arrangement immediately following seeding. In contrast, on day 2 the spheroids appeared less diffuse (Fig. 2G) and were flatter at the top (Fig. 2L). As the cells proliferated and the spheroids became larger, internal gaps formed within the spheroid bulk by day 6 (Fig. 2H), although these gaps were not visible externally (Fig. 2M). Despite the presence of internal gaps, attenuation due to spheroid density reduced the OCT signal from deeper regions (Fig. 2H). After 10 and 14 days, culture media contained many suspended cells, and volumetric images revealed individual cells along the slope of the gel (Fig. 2N,O).

The average spheroid volume, internal gap volume, external surface area, and sphericity of live and fixed spheroids over 14 days of growth are shown in Figure 3. Generally, we observed similar morphological trends in both live and fixed spheroids. The spheroid volume, internal gap volume, and external surface area of live spheroids all initially decreased following cell seeding, reached a minimum on day 4, then increased and repeated this trend twice more (Fig. 3A-C). However, sphericity continued to increase until it peaked on day 7 (Fig. 3D).

**Figure 3.**
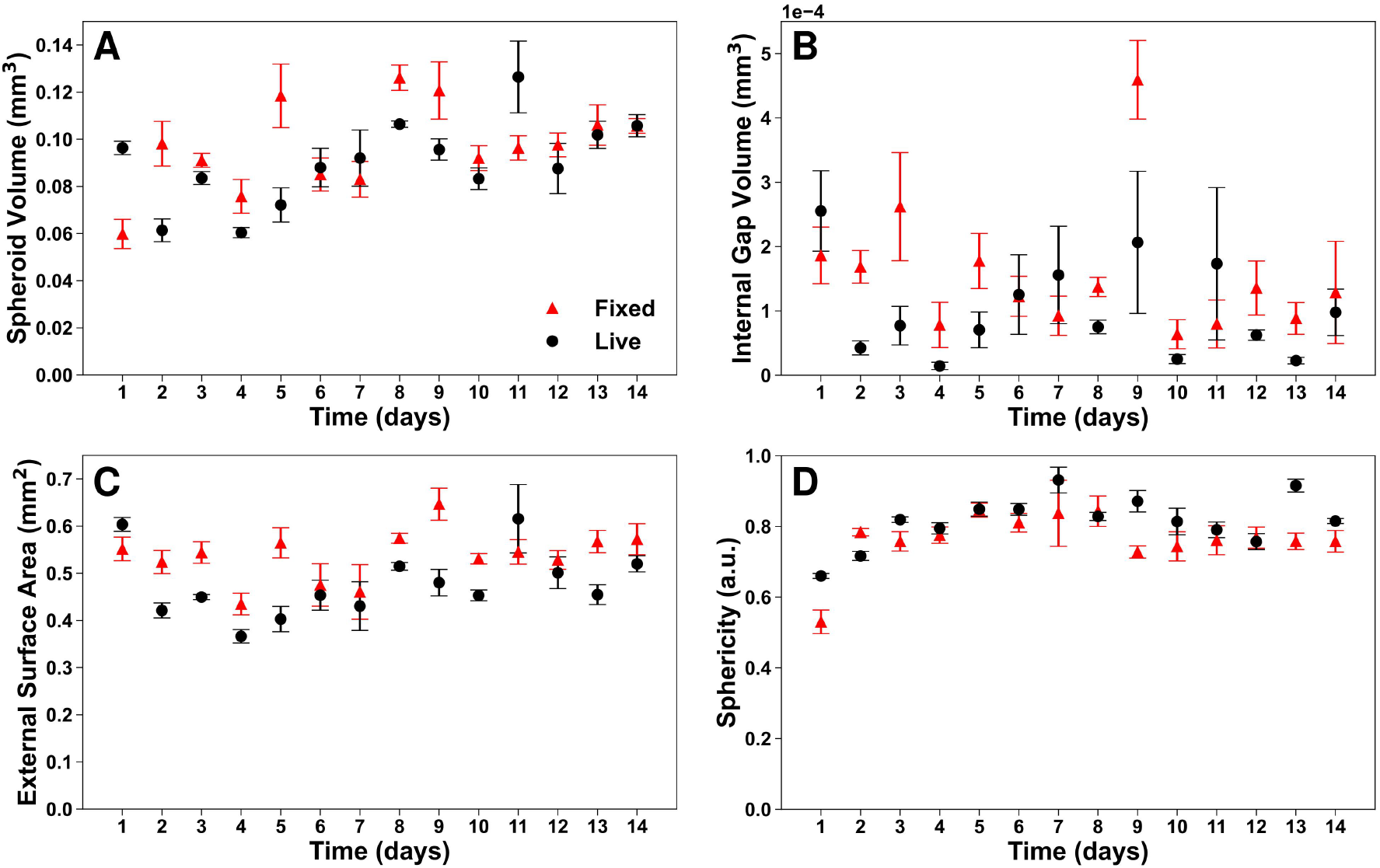
Numerical morphological analysis of masked volumetric OCT images of prostate tumor spheroids over 14 days of growth. (A) spheroid volume, (B) internal gap volume, (C) external surface area, and (D) sphericity given by Eq. 2. Average measurement of live (black) and fixed (red) spheroids are presented with standard deviation error bar.

### 3.2 Dynamic OCT

Figure 4 compares OCT, dOCT, and FM enface images of spheroids on days 1, 2, 6, 10, and 14. The first and second rows respectively display OCT and corresponding dOCT maximum projection enface images of the same spheroids shown in Figure 2, while the third row shows FM enface images of separately imaged spheroids.

**Figure 4.**
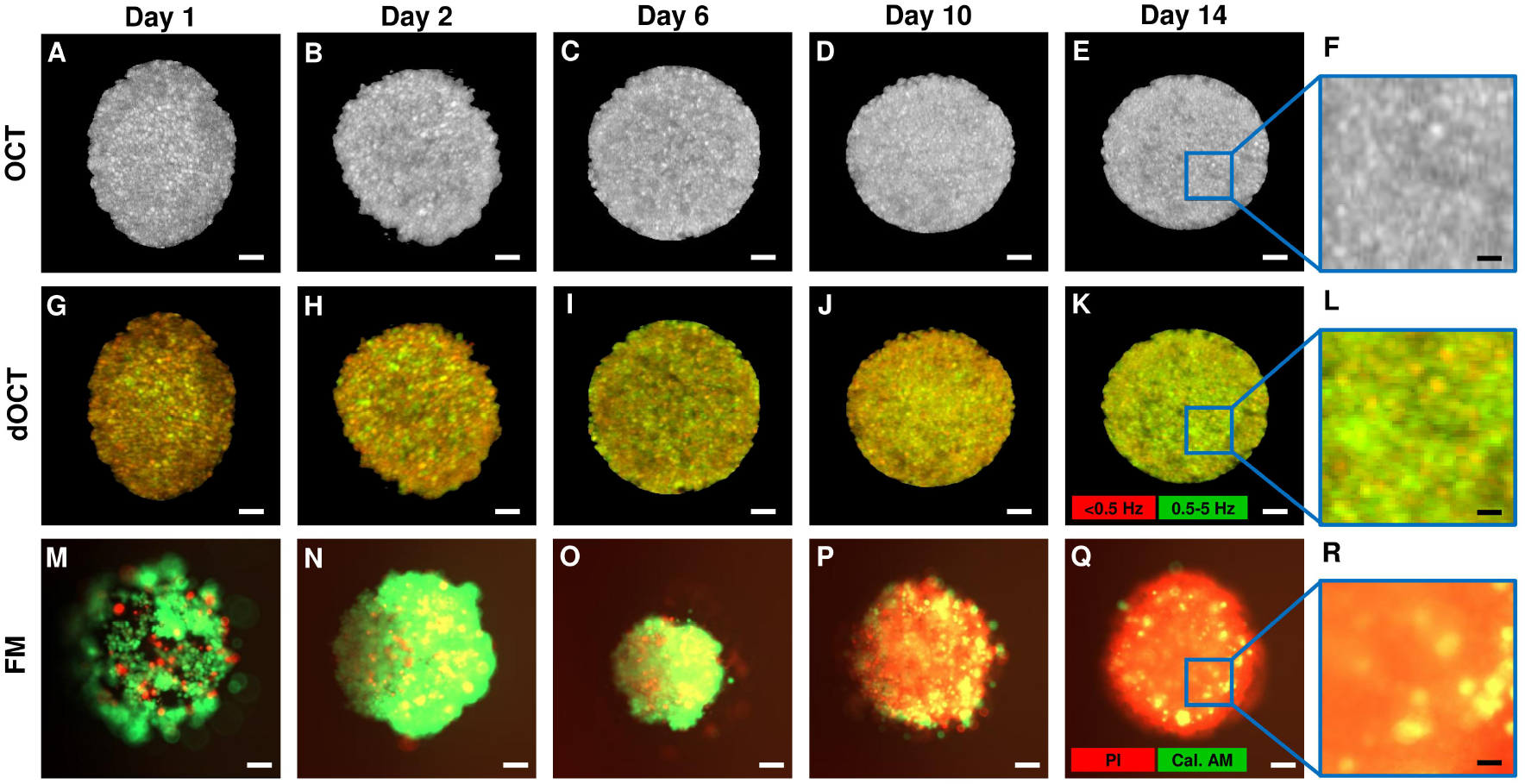
Maximum projection OCT, maximum projection dOCT, and FM enface images of prostate tumor spheroids 1, 2, 6, 10, and 14 days after cell seeding. Blue inset in last column shows 4X magnification with 25 *µ*m scale bars. All other scale bars are 100 *µ*m.

While the OCT enface images resolved individual cells and did not vary significantly over 14 days, the dOCT enface images resolved cells equally well while also indicating dynamic changes over time. Overall, the dOCT green channel intensity increased throughout the 14-day period. On day 1, the dOCT red channel intensity was higher at the spheroid periphery, whereas the green channel exhibited higher intensity in the central region (Fig. 4G). By day 2, the dOCT green channel intensity increased and extended to cover a larger area, although the red channel intensity remained higher along the periphery (Fig. 4H). On days 6, 10, and 14, both the dOCT red and green channel intensities were more uniformly dispersed across the enface image (Fig. 4I-K).

In contrast, on day 1, FM showed predominantly live (green channel) intensity at the spheroid periphery, some central red channel intensity indicating dead cells, and dark regions suggesting a diffuse cellular arrangement (Fig. 4M). By day 2, the spheroid had condensed, and a central yellow hue emerged due to the overlap of green and red channel intensities (Fig. 4N). Subsequently, the coverage of the central red channel intensity expanded on day 6 (Fig. 4O), filled the spheroid’s cross-sectional area by day 10 (Fig. 4P), and overwhelmed the green channel intensity by day 14 (Fig. 4Q).

The last column of Figure 4 shows 4× enlarged cutouts of each day 14 enface image, demonstrating the improved resolution of the OCT system compared to the fluorescent microscope. Notably, localized regions of high intensity in the OCT enface (Fig. 4F) correspond to either the green or red channel intensities in the dOCT enface (Fig. 4L).

Figure 5 summarizes the numerical dOCT analysis. The raw dynamic signal of live and fixed spheroids, as well as the normalized dynamic signal of live spheroids over 14 days of growth, are shown in Figure 5A and B, respectively. The top row of Figure 5C shows volumetric and enface dOCT images of a live spheroid, and the bottom row shows volumetric and enface dOCT images of the same spheroid immediately after it was fixed for one hour with 4% formaldehyde. The raw dynamic signal of live spheroids remained consistently larger than that of fixed spheroids, however the raw dynamic signal of fixed spheroids was not negligible, and both signals increased gradually over 14 days (Fig. 5A). Correspondingly, although the spheroid appears more red after one hour of fixing, some cells in the fixed spheroid still appear intensely green (Fig. 5C). Figure 5D shows the normalized AB absorbance over 14 days of growth. After normalization, the dynamic signal and AB absorbance of live spheroids both initially increased, decreased on day 5, and then increased and decreased once more (Fig. 5B,D).

**Figure 5.**
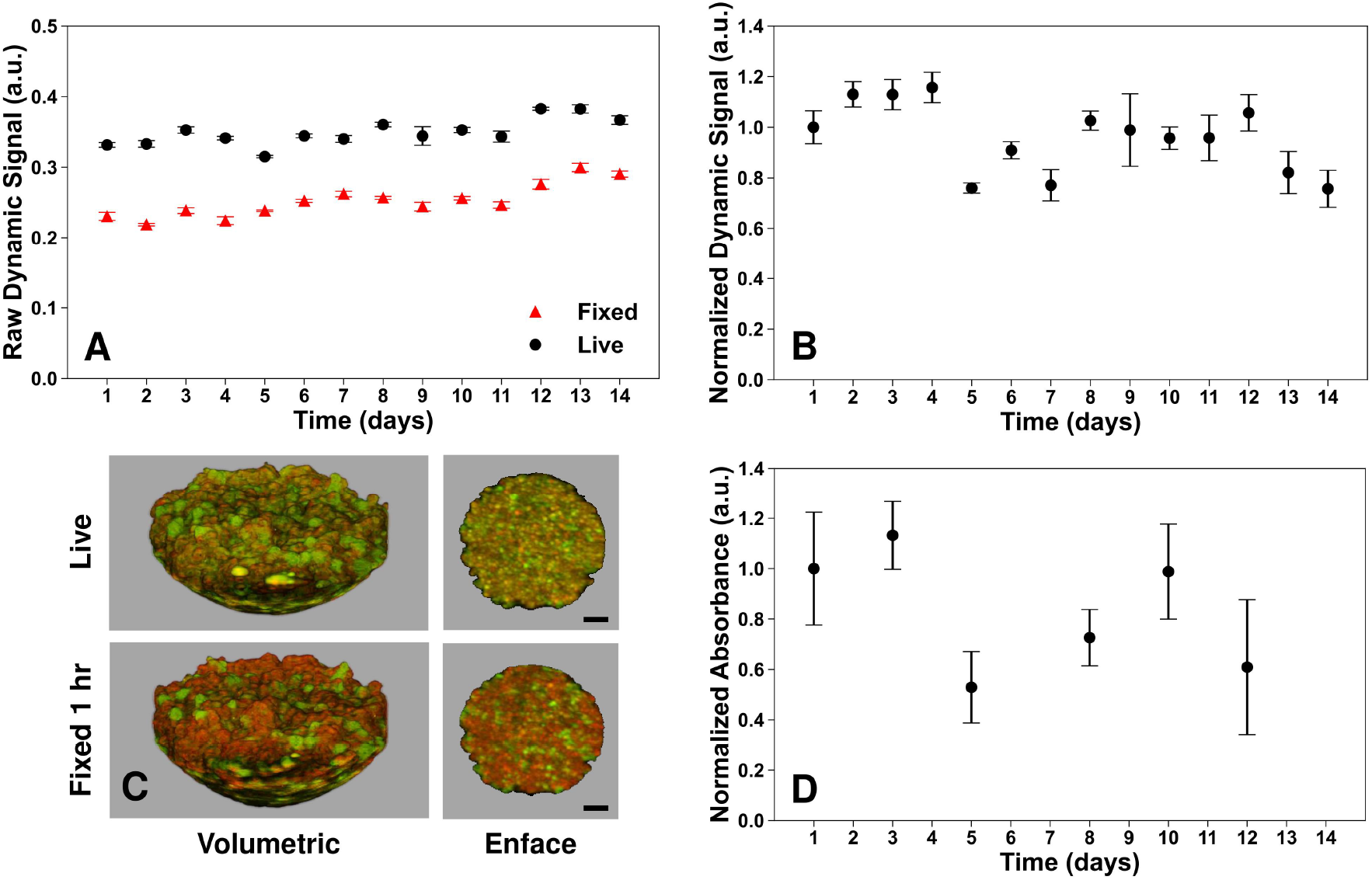
Numerical dynamic analysis of masked volumetric dOCT images of prostate tumor spheroids over 14 days of growth. (A) raw dynamic signal of live (black) and fixed (red) spheroids given by Eq. 3. (B) normalized dynamic signal given by Eq. 4. (C) Volumetric and maximum projection enface images of a live spheroid and the same spheroid after 1 hour of fixation with 4% formaldehyde. (D) Normalized absorbance of AB proliferation assay. Average measurements are presented with standard deviation error bar. All scale bars are 100 *µ*m.

## 4 Discussion

### 4.1 Spheroid seeding and morphology

While 3D tumor spheroids are widely recognized as a more physiological *in vitro* model of tumor behavior than 2D monolayer cultures [12, 13], the “3D Petri Dish” is not a conventional technique for spheroid seeding and cultivation. Comparison between various spheroid formation methods is complicated by the influence of both cell-to-cell adherence, which varies by cell type, and the 3D culture environment on cellular arrangement [71, 72]. Previous studies have used OCT to investigate tumor spheroids formed in low-attachment well-plates [24, 30–33, 36,38], by hanging-drop [21], in microfluidic systems [28], or embedded in a collagen or gel matrix [27, 29, 34, 35]. The PC3 cells employed in our study adhere loosely [71] and preferentially to neighboring cells over the agarose gel, as demonstrated by the cell clumping observed within 5 minutes of seeding (Fig. 2B). After cell seeding, we observed that initially diffuse spheroids condensed before their volume generally increased, albeit with some fluctuations (Fig. 3A). Similar trends were noted by Sharma et al. [21] and Tartagni et al. [73], though without voxel-wise masking that characterized internal gaps. While we observed that sphericity peaked at nearly 1 on day 7 (Fig. 3D), the spheroids did not appear “spherical” in most volumetric images (Fig. 2K-O), a finding consistent with previous visual [21] and numerical observations [34].

### 4.2 Cyclical proliferation

We observed similar trends in the dynamic signal and AB absorbance after normalization. Both metrics increased approximately 15% from the initial day 1 measurement by day 3 (Fig. 5B, D). Despite the AB proliferation assay suggesting an increase in cell number over this period, on day 4, the volume and external surface area of live spheroids had decreased to approximately two-thirds of their initial day 1 values (Fig. 3A, C). However, the internal gap volume similarly decreased and reached a minimum on day 4 (Fig. 3B), suggesting that the initially large volume and surface area in newly seeded spheroids arose from dispersed cells that subsequently condensed into more tightly aggregated structures. After this compaction, the spheroid volume, internal gap volume, and external surface area all increased (Fig. 3A-C). We propose that once cells became tightly aggregated, some died due to insufficient nutrition and detached from their neighbors. As a result, the spheroid partially dispersed, allowing central cells better access to nutrients and thus improved growth. This interpretation is supported by the normalized dynamic signal and AB absorbance, which both reached a minimum on day 5, then increased once again (Fig. 5B, D). The subsequent period of mostly increasing spheroid volume, internal gap volume, external surface area, and normalized dynamic signal continued until approximately day 8. Then, the pattern repeated: the three morphological metrics decreased until approximately day 10 before increasing on day 11 (Fig. 3A-C). Lagging by a day, the normalized dynamic signal decreased until day 11 before increasing on day 12 (Fig. 5B). Although less frequent measurements complicate direct comparison, normalized AB absorbance also rose again before subsequently decreasing (Fig. 5D). The increase in spheroid volume, internal gap volume, and external surface area on day 14 (Fig. 3A-C) indicates cell death and detachment within the spheroids, which is further supported by the low normalized dynamic signal (Fig. 5B) and high FM red channel intensity (Fig. 4Q) observed on day 14.

### 4.3 Necrotic spheroid core

FM revealed a predominantly green spheroid periphery with central red intensity that increased over time, suggesting the formation of a necrotic core of dead cells surrounded by proliferating live cells (Fig. 4M-Q). This is an established phenomenon previously observed both by conventional biological methods [12, 13] and dOCT [30, 31]. In contrast, dOCT images showed increasing green channel intensity throughout the 14-day period (Fig. 4G-K) and the raw dynamic signal measured via dOCT in live spheroids did not follow the trend observed with AB (Fig. 5B,D). Notably, the raw dynamic signal in spheroids killed by formaldehyde fixation also increased throughout the experiment (Fig. 5A). We assume that the dynamic motion observed in fixed spheroids did not change over the two-week experimental period because an identical fixing procedure was followed each day. Therefore, the variation in fixed dynamic signal over time suggests that the system’s performance or the influence of factors unrelated to cellular metabolism also varied with time. To correct for these factors external to the sample, the dynamic signal of fixed spheroids was subtracted as background from live spheroids, and the subsequent normalized dynamic signal measured via dOCT agreed well numerically with AB proliferation assay (Fig. 5B,D). However, dOCT images visually represent the raw dynamic signal before averaging and subtraction of the corresponding fixed measurement. Since spheroids were maintained at 4°C for approximately 24 hours post-fixation, the dOCT images of fixed spheroids were not registered to their live counterparts. Consequently, the dOCT images could not be similarly fixation-subtracted for an improved qualitative comparison to FM.

### 4.4 Origin of dynamic signal

We observed that dOCT images of live spheroids were significantly more green than fixed spheroids (Fig. 5A). As fixation eliminates metabolic activity by killing cells without damaging the spheroid structure, this differentiation suggests that the LF-dOCT system could distinguish temporal intensity fluctuations due to cellular metabolism. Furthermore, the average dynamic signal throughout the spheroid bulk, once fixation-subtracted and normalized, demonstrated excellent agreement with AB (Fig. 5B), a well-established indicator of cellular metabolism [17]. However, the longitudinal development of a necrotic spheroid core revealed by FM (Fig. 4M-Q) was not spatially well-resolved in dOCT images, which instead showed higher green intensity in the spheroid core (Fig. 4G-K). While AB is specific to cellular activity and becomes fluorescent through reduction by cytoplasmic and mitochondrial enzymes [74–76], the sub-cellular motion measured by dOCT may also arise from physical changes in cell morphology and volume. Cells undergoing necrosis exhibit physical changes, such as cell swelling, nuclear disintegration called karyolysis, and ultimately degradation of the cellular membrane and leakage of cellular contents [77, 78]. Following necrosis, these cellular fragments may have been subject to Brownian motion and further contributed to the high frequency motion detected in the spheroid core.

Unlike necrotic cell death due to competition for resources in a confined space, formaldehyde fixation kills cells quickly and efficiently, preserving spheroid structure by cross-linking proteins through a process completed within 24 to 48 hours [79]. Fixed spheroids were maintained at 4°C for approximately 24 hours to reduce motion caused by the cross-linking process. Nevertheless, some cells throughout the spheroid bulk remained intensely green in dOCT images after fixation (Fig. 5C). While some of these cells may have survived the fixing process, the localized high frequency motion measured by dOCT in fixed spheroids could also be due to non-metabolic motion. Although cells killed by fixation are expected to remain tightly adhered to neighbors, some cells may have died by necrosis before fixation and the resulting cell bodies and fragments may have experienced Brownian motion.

### 4.5 LF-dOCT system and dynamic analysis methods

The current system achieves a sensitivity of 93 dB, using a 2,000 fps camera speed and 3.5 mW of incident power at the imaging plane. This represents approximately a 3 dB improvement in sensitivity compared to the initial report of the dOCT platform [37], which can be attributed to finer alignments. Our imaging protocol acquired each sub-volume at a rate of 10.7 Hz. By the Nyquist-Shannon sampling theorem [80], the fastest dynamics observable to our method occurred at just over 5 Hz: the maximum threshold of our green frequency band. While our system and method appear sufficient for numerically accurate extraction of spatially-averaged and fixation-subtracted dynamic signals (Fig. 5B,D), the spheroid’s necrotic core is not visually resolved (Fig. 4). The following section discusses various system characteristics that may have contributed.

#### 4.5.1 Temporal resolution

To observe cellular activity with dOCT, there must be sufficient temporal resolution of the OCT signal. Dynamic contrast is provided by ATP-consuming processes and the active movement of organelles [26], which are estimated to be in the range of 0.1 to 20 Hz [81, 82]. By detecting frequencies up to 28 Hz, Wang et al. [38] visually resolved the necrotic spheroid core. They generated HSB images using the mean frequency, inverse frequency bandwidth, and running standard deviation technique developed by Scholler et al. [42]. The frequency banding method we employed similarly analyzes the OCT spectrum and has been used to detect frequencies up to 12 Hz [55], 20 Hz [58], or 25 HZ [54, 56, 57]. Nevertheless, Liu et al. [67] used mean frequencies limited to the range of 0.16 Hz to 1 Hz to distinguish live and dead cells in mouse allograft tumors. In fact, the first reported dOCT study differentiated between viable, necrotic, and apoptotic human fibroblasts with single B-scans acquired every few minutes [52]. Consequently, while our maximum detectable frequency of 5 Hz may have contributed to the inability to resolve the necrotic spheroid core, it was likely not the most significant factor.

#### 4.5.2 Sensitivity

The limited sensitivity of our LF-dOCT platform may have also contributed to the undistinguished necrotic core. Because dOCT analysis relies on OCT intensity fluctuations, a higher system sensitivity should allow for more accurate extraction of dynamic signals. In general, acquisition speed is increased at the cost of reduced sensitivity [83]. The maximum frequency thresholds of 12 Hz [55], and 25 HZ [56, 57] discussed in section 4.5.1 were achieved with a PSc-OCT system that demonstrated a sensitivity of 51 dB at a depth of 80 *µ*m. Alternatively, the studies by El-Sadek et al. [30–33] successfully resolved the necrotic core in human breast adenocarcinoma (MCF-7) spheroids by employing a PSc-OCT system with a sensitivity of 104 dB. Compared to our system, their 11 dB higher system sensitivity enabled them to directly evaluate OCT intensity fluctuations via the LIV and OCDS metrics. Likewise, Tan et al. [36] implemented MAV, another direct variation method, to investigate MCF-7 spheroids with a commercial PSc-OCT system. However, akin to our results, they did not observe a necrotic spheroid core and demonstrated quantitative agreement with viability measured by Trypan Blue, a biological method similar to AB that provides an average measurement across many spheroids.

#### 4.5.3 Frequency band thresholds

In contrast to these methods, to mitigate the strong influence of noise inherent in direct variation methods, we addressed our reduced sensitivity by employing the frequency banding technique introduced by Apelian et al. [61]. By integrating the spectrum of OCT intensity fluctuations, this approach effectively averages over a range of signals, thereby diminishing the impact of random noise. Previous work has qualitatively compared this technique to fluorescence imaging [62] and histological staining [54, 56–58] to differentiate various cellular and intracellular morphological structures. However, to the best of our knowledge, the work we describe here is the first to quantitatively validate it to an established biological method. Meanwhile, the HSB spectrum-based method [42] has been used to assess the viability of HeLA cervical cancer cells cultured in 2D monolayers and qualitatively compared with Trypan Blue, which stains cells with broken cell membranes [65]. The mean frequency computed in this method has also been quantitatively validated using FM images captured by confocal laser scanning microscopy [38] or multiphoton microscopy with registration to the OCT equivalent [67]. Since Wang et al. [38] successfully resolved the necrotic spheroid core with a spectrum-based dOCT method, perhaps the thresholds imposed by our frequency banding approach contributed to our undistinguished necrotic core. Future studies should quantitatively investigate how differences in system sensitivity, the threshold between the red and green frequency bands, and the removal of a blue frequency band impact the accuracy of dOCT signal extraction.

### 4.6 Reflection artifacts and pseudo-longitudinal protocol

As is common in OCT systems, reflection artifacts were prevalent and difficult to mitigate. This issue is particularly pronounced in LF-OCT systems due to the absence of confocal gating that is leveraged by PSc-OCT systems. While imaging the seeding process, the sample was not adjusted; however, an artifact that was not present at 10 minutes after seeding (Fig. 2D) appeared at 60 minutes (Fig. 2D, red arrow). The sudden appearance of the artifact was likely due to the evaporation of the culture medium over the course of imaging, which altered the angle of the liquid surface. Reflection artifacts were minimized in regions of the gel where the liquid surface was slightly curved due to surface tension, resulting in fewer direct reflections reaching the camera. Consequently, spheroids located in these edge areas were selected more frequently for imaging. However, location within the 9 × 9 well array inside the gel influences spheroid size, as seen in the yellow inset in Figure 1B. Although 15 live and fixed spheroids were imaged each day to form a representative sample, this selection process likely introduced some bias into the morphological measurements. Future work is encouraged to record spheroid location and identify its statistical impact.

Additionally, the Petri dish’s top cover introduced significant reflection artifacts and was therefore removed during imaging, potentially exposing the cell culture to contamination. As a result, samples were not returned to the sterile incubator and we employed a suboptimal pseudo-longitudinal protocol that imaged different spheroids over the 14-day growth period. To conduct a truly non-invasive longitudinal study, imaging should be performed through the Petri dish cover, or more ideally, with an incubator integrated into the OCT imaging stage. Such integration would not only prevent the need for spheroid transportation – a mechanically stressful process that may alter cellular arrangement – but also enable the growth of individual spheroids to be monitored over time. Even in this integrated scenario, we suggest including a subset of spheroids that are fixed and imaged during each live imaging session. This would create a relevant metabolic control and establish a baseline of system performance for comparison. Numerical analysis of our pseudo-longitudinal protocol, which fixed spheroids at the time of each live imaging session, revealed significant changes in fixed spheroids over time (Fig. 3). These changes may influence the dynamic response to non-metabolic motion. A limitation of our study is that, since each nominal day of fixed spheroid measurement was acquired the following day, fixed subtraction in the normalized dynamic signal does not precisely account for the system performance on the exact day of live spheroid imaging. However, because imaging occurred every 24 hours, the system performance is not expected to undergo significant changes within that time frame.

### 4.7 Signal attenuation

Another factor complicating the interpretation of dOCT results was attenuation caused by spheroid condensation and growth beyond the system’s 260 *µ*m depth of focus (DOF) in conjunction with the signal-to-noise ratio (SNR) roll-off inherent to the spectral domain dOCT platform. Although intensity was notably reduced in the lower portions of large spheroids in the morphological images (Fig. 2H), the spheroid masks were manually extended to include these attenuated regions in the morphological analysis. However, the dynamic signals could no longer be accurately extracted, and we excluded the low-intensity regions of the dOCT data using the intensity filter described in Methods Section 2.4.2. To address this issue, several approaches could be considered: reducing the lateral resolution further to increase the DOF; switching from a spectral domain to a swept-source OCT system to avoid SNR roll-off; or using a longer central wavelength light source, such as 1310 nm, to enhance penetration depth and improve overall performance.

## 5 Conclusion

In conclusion, we acquired volumetric dOCT images of prostate tumor spheroids with a spectral-domain LF-dOCT system to characterize structural morphology and cellular dynamics over two weeks of growth. By comparing live spheroids with those killed by formaldehyde fixation, we confirmed that the system’s sensitivity was sufficient to detect dynamic signal variations associated with cellular metabolism. Numerical analysis of dOCT images was in excellent agreement with results from AB proliferation assay. In future work, we plan to further improve the system’s sensitivity to enhance the reliability of the dOCT platform and to incorporate an incubator for truly non-invasive, repeatable cellular dOCT measurements.

## Funding

Canadian Institutes of Health Research (202104PJT-461005); Natural Sciences and Engineering Research Council of Canada (RTI-2021-00780, RTI-2022-00169); Mitacs (53162-10628); Prostate Cancer Fight Foundation.

## Acknowledgments

It is with sadness that the authors acknowledge the recent passing of the senior author, Dr. Kostadinka Bizheva. The authors would like to gratefully acknowledge Dr. Mohammad Kohandel for generous use of his laboratory equipment. We thank Dr. Brian Ingalls and Atiyeh Ahmadi for their expertise and use of their fluorescent microscope, and Dr. Qing-Bin Lu and Olya Changizi for access and assistance with their microplate reader. Lastly, we are grateful for the help of Catherine McKenna for assisting in cell culture preparation.

## Disclosures

The authors declare no conflicts of interest.

## Data availability

Data underlying the results presented in this paper are not publicly available at this time but may be obtained from the authors upon reasonable request.

## Code availability

The code used to process the OCT and dOCT images is publicly available on GitHub at github.com/skswanso/D-OCT.git

## Notes

### Competing Interest Statement

The authors have declared no competing interest.

